# Barley ROP-INTERACTIVE PARTNER-a organizes into RAC1- and MICROTUBULE-ASSOCIATED ROP-GTPASE ACTIVATING PROTEIN 1-dependent membrane domains

**DOI:** 10.1101/693804

**Authors:** Caroline Hoefle, Christopher McCollum, Ralph Hückelhoven

**Affiliations:** Chair of Phytopathology, TUM School of Life Sciences Weihenstephan, Technical University of Munich, Emil Ramann Str. 2, 85354 Freising, Germany

**Keywords:** *Arabidopsis thaliana*, *Hordeum vulgare*, interactor of constitutive active ROPs, membrane asymmetry, microtubule, RAC GTPase, ROP GTPase, susceptibility, resistance

## Abstract

Small ROP (also called RAC) GTPases are key factors in polar cell development and in interaction with the environment. ROP-Interactive Partner (RIP) proteins are predicted scaffold or ROP-effector proteins, which function downstream of activated GTP-loaded ROP proteins in establishing membrane heterogeneity and cellular organization. Grass ROP proteins function in cell polarity, resistance and susceptibility to fungal pathogens but grass RIP proteins are little understood.

We found that the barley (*Hordeum vulgare* L.) RIPa protein can interact with barley ROPs in yeast. Fluorescent-tagged RIPa, when co-expressed with the constitutively activated ROP protein CA RAC1, accumulates at the cell periphery or plasma membrane. Additionally, RIPa, locates into membrane domains, which are laterally restricted by microtubules, when co-expressed with RAC1 and MICROTUBULE-ASSOCIATED ROP-GTPASE ACTIVATING PROTEIN 1. Both structural integrity of MICROTUBULE-ASSOCIATED ROP-GTPASE ACTIVATING PROTEIN 1 and microtubule stability are key to maintenance of RIPa-labeled membrane domains. In this context, RIPa also accumulates at the interface of barley and invading hyphae of the powdery mildew fungus *Blumeria graminis* f.sp. *hordei*.

Data suggest that barley RIPa interacts with barley ROPs and specifies RAC1 activity-associated membrane domains with potential signaling capacity. Lateral diffusion of this RAC1 signaling capacity is restricted the resulting membrane heterogeneity requires intact microtubules and MICROTUBULE-ASSOCIATED ROP-GTPASE ACTIVATING PROTEIN 1. Focal accumulation of RIPa at sites of fungal attack may indicate locally restricted ROP activity at sites of fungal invasion.

## Introduction

In plants, ROP (RHO of plants) small GTPases are the only members of the RHO protein family, which consists of several subfamilies (RHO, RAC, CDC42, Rnd und RhoBTB) in mammals [1, 2]. ROPs organize a bunch of cellular processes as signaling GTPase. Among the most prominent ROP-regulated events are the subcellular organization of the cytoskeleton and vesicular traffic [3]. ROP-regulated cellular organization is crucial for normal plant development e.g. in polar cell growth or asymmetric cell division but also in interaction with the environment e.g. in regulation of stomata aperture or in interaction with pathogens. ROP activity is tightly regulated via proteins that facilitate hydrolysis and exchange of ROP-bound nucleotides. ROP-GDP is the signaling-inactive form of ROP and can be further controlled by ROP-GDIs (ROP-guanine nucleotide dissociation inhibitors) that bind to ROP-GDP. ROP-GDIs support cytosolic localization of ROPs most likely by direct binding of isoprenyl-residues at the C-terminus of type I ROPs, which carry a CAAX-box prenylation motif. ROP-GDP further can interact with different types of ROP guanine nucleotide exchange factors (GEFs), which support the release of GDP and binding of GTP. This turns the protein into activated ROP-GTP that signals downstream. ROP GTPase-activating proteins (GAPs) then can switch off activated ROPs again by supporting the otherwise low intrinsic GTPase function of ROPs and facilitating GTP hydrolysis [3, 4]. Negatively charged lipids at the inner leaflet of the plasma membrane may further function in ROP-positioning and signaling [5, 6].

In barley, distinct ROP GTPases are susceptibility factors in the interaction with the powdery mildew fungus *Blumeria graminis* f.sp. *hordei* (*Bgh*). Several ROPs, when constitutively activated (CA) by mutations in the GTPase domain, can support invasion of epidermal cells by fungal hyphae, which subsequently form a haustorium as a feeding cell in a living epidermal cell of barley [7]. *Vice versa*, sequence-specific RNA interference for silencing *RACB* renders barley less susceptible to fungal invasion and limits disease development [8, 9]. RACB’s physiological function is described in polar cell development during formation of root hairs and leaf stomata complexes [10]. Since *Bgh* appears to target RACB directly by an virulence effector, it was suggested that the fungus exploits a plant polar cell developmental pathway for the accommodation of haustoria in living barley cells [11]. Another barley ROP called RAC1, has a less well understood function in the interaction with *Bgh*. Transient expression of CA RAC1 in single epidermal cells did not render barley supersusceptible [7]. However, the same open reading frame, when stably expressed in transgenic barley, supported fungal penetration but also the generation of reactive oxygen species in non-penetrated cells. CA RAC1 further supported barley resistance to the rice blast fungus *Magnaporthe oryzae*, similar to what was reported before for the function of rice RAC1, which is 86% identical to barley RAC1 [4, 12].

The barley genome encodes several predicted ROP-GAP proteins, but only the MAGAP1 (MICROTUBULE-ASSOCIATED ROP-GTPASE ACTIVATING PROTEIN 1) has been characterized thus far. MAGAP1 contains a CRIB motif (for <u>C</u>DC42/<u>R</u>AC-Interactive <u>B</u>inding) and can bind to both RACB and RAC1 and is associated with microtubules. However, besides a localization at MTs, MAGAP1 positions at the cell periphery when recruited by CA RACB and to a minor extent in the cytoplasm. MAGAP1 is considered as a functional antagonist of RACB because MAGAP1 overexpression limits susceptibility whereas *MAGAP1* silencing supports susceptibility to penetration by *Bgh* [9]. Additionally, potentially ROP-regulated stability and polarity of MTs is associated with resistance to fungal penetration in barley [9, 11, 13].

ROP-GTP signals downstream via protein-protein interaction that depends of the ROP-loaded nucleotide and hence the three-dimensional constitution of ROPs. Proteins, which mediate ROP downstream effects, are commonly called ROP-effectors. However, not all ROP-effectors directly fulfill a function in cellular organization but instead are suggested to be scaffolds or adapter proteins that link activated ROPs with downstream factors. RIPs (<u>R</u>OP-Interactive <u>P</u>artner, also called Interactor of Constitutive Active ROPs [ICR]) and RICs (<u>R</u>OP-Interactive <u>C</u>RIB motif-containing proteins) are such ROP-effectors without known biochemical but potential ROP-scaffolding function [3].

## Results

### Barley RIPa is a ROP binding protein

Because ROP signaling and microtubule organization seems to be important in interaction of barley and *Bgh*, we looked for candidate proteins that potentially are involved in both processes. *Arabidopsis thaliana* RIP3 (also called ICR5 and microtubule depletion domain 1, MIDD1) can interact with ROPs and MT-associated kinesin13A *in planta* [14]. Oda and co-workers found RIP3/MIDD1 to be part of a ROP regulatory module, which determines MT organization and subcellular cell wall deposition in xylem cells [15-17]. We therefore speculated that barley proteins with homology to RIP3 (AT3G53350) can act in ROP signaling during fungal invasion or defensive plant cell wall apposition (see also [18]). The barley locus HORVU3Hr1G087430.11 (protein accession F2DI37_HORVV) encodes the barley protein with the highest similarity to Arabidopsis RIP3). However, protein identity between these Arabidopsis and barley RIP proteins is only 36% and the barley protein is with 510 amino acids much longer than Arabidopsis RIP3 with 396 amino acids. We thus named the barley protein RIPa instead of RIP3 because based on that we cannot predict whether barley RIPa is indeed the orthologue of Arabidopsis RIP3. To confirm that RIPa might be a ROP-binding protein, we checked protein-protein interaction in a targeted yeast-two-hybrid assay and found that RIPa interacts with RACB and RAC1 from barley as well as with CA versions of these proteins but not with dominant negative versions (Fig. 1). RIPa appears thus to be able to interact in yeast with so-called type I ROPs carrying a carboxyterminal CAAX-box prenylation signal as well as with type II ROPs that are predicted to be constitutively palmitoylated [7, 19].

**Fig. 1:**
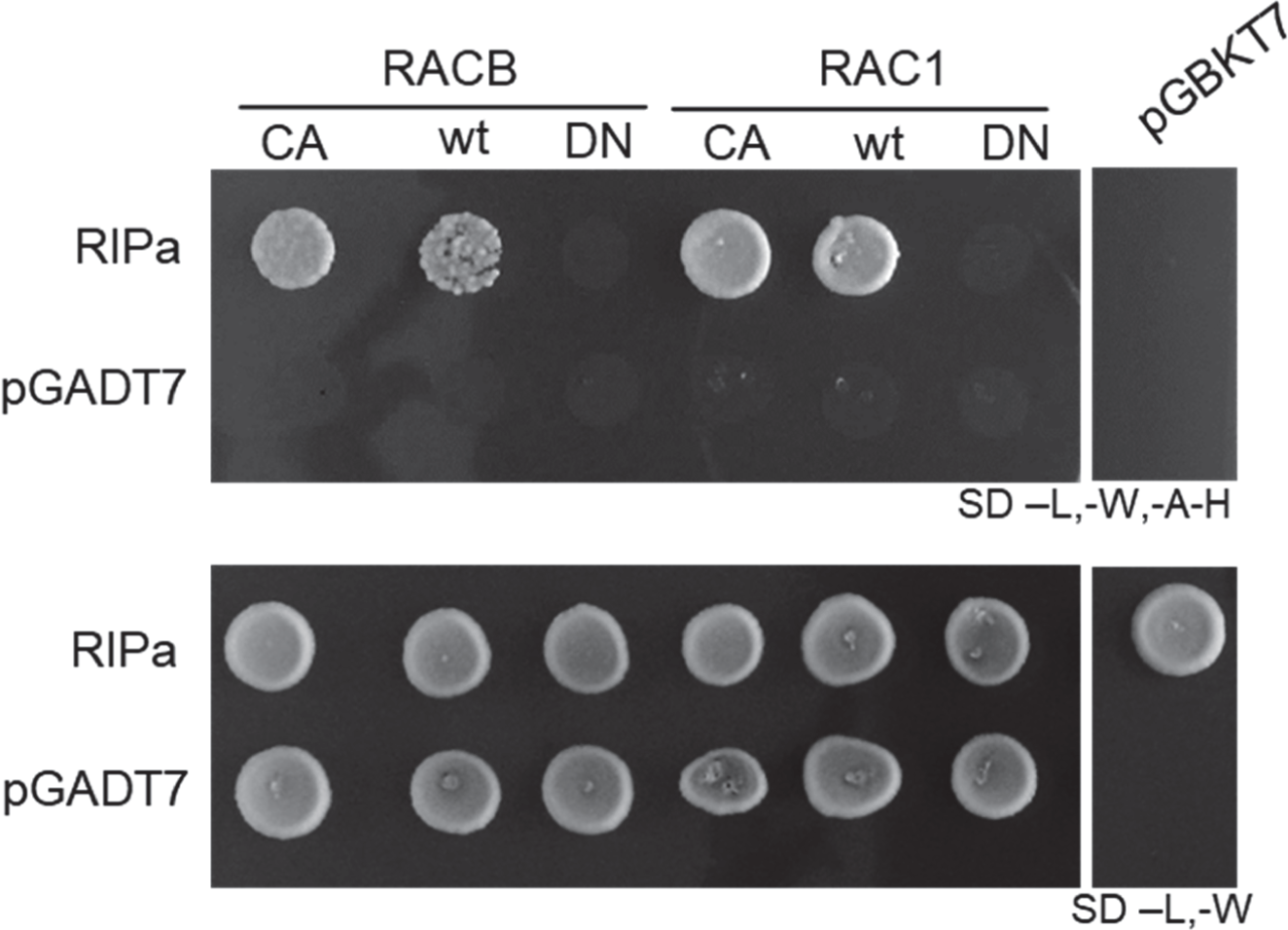
Barley RIPa interacts with barley type I and type II ROPs in yeast. Bait- and prey-construct transformed yeast cells were dropped on either transformation-selected (SD -L-W) or interaction-selective (SD –L,-W,-A-H) medium. pGADT7 and pGBKT7 present empty vector controls to exclude auto-activity of respective ROP or RIPa constructs.

### ROPs can influence subcellular localization of RIPa

We then studied subcellular localization of RIPa by confocal laser scanning microscopy. When we expressed a yellow fluorescing fusion protein, YFP-RIPa, the fluorescence signal was always detectable in the cytoplasm and strong in undefined speckels, which were little mobile and only co-localized partially with the microtubule (MT)-marker RFP-MAGAP1-Cterm, which contains the MT-binding domain of MAGAP1 but does not interact with ROPs because it lacks the ROP-binding CRIB and GAP domains (see below and Hoefle, 2011 #398) (Fig. 2).

**Fig. 2:**
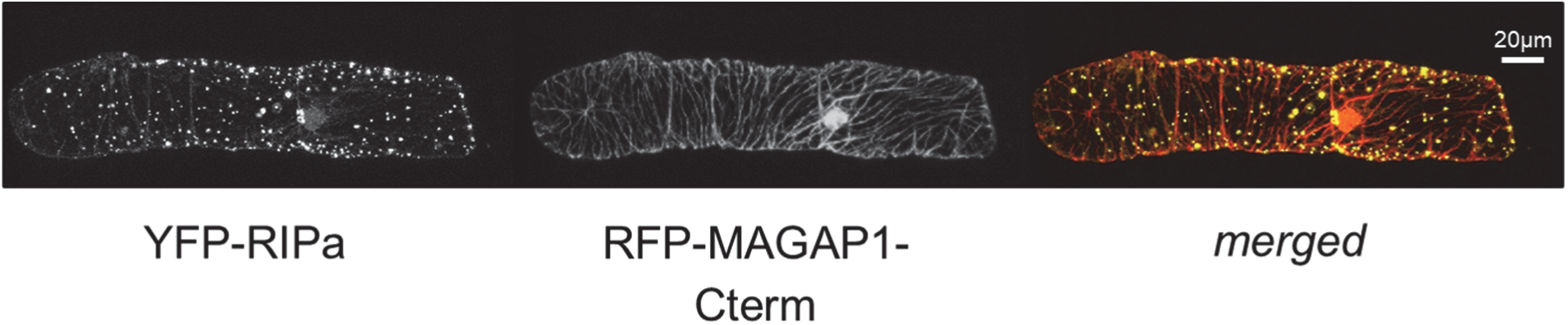
Single cell-expressed barley YFP-RIPa localizes to immobile speckles and the cytoplasm. Whole cell Z-stack images were taken 24 h after biolistic transformation of barley epidermal cells. The MT-marker RFP-MAGAP1-Cterm was co-expressed to visualize MTs.

We hypothesized that the speckled localization of YFP-RIPa represents protein aggregates that form when a scaffold protein is expressed without a corresponding amount of protein binding partners. RIPa could also interact with itself in yeast-2-hybid assays and hence might form multimers when ectopically expressed (Additional file 1). To test, whether co-expression of potential binding partners might change subcellular localization of YFP-RIPa, we co-expressed RAC1, CA RAC1 and DN RAC1. Astonishingly, both expression of RAC1 or CA RAC1 completely changed subcellular localization of YFP-RIPa. RAC1 fully recruited YFP-RIPa to the cell periphery or plasma membrane and to a minor extent also to MTs, whereas CA RAC1 recruited YFP-RIPa exclusively to the cell periphery/plasma membrane. DN RAC1 did not recuit YFP-RIPa or perhaps even enhanced protein aggregation in speckles (Figure 3). Together, data suggest that CA or wildtype switchable RAC1 can influence the localization of YFP-RIPa most likely by direct protein interaction. In figure 3, a red fluorescing MT-marker was co-expressed. To further exclude that the marker influenced YFP-RIPa localization, we repeated the experiments with free mCherry as cytoplasmic and nucleleoplasmic marker. Similar to was was observed before, CA RAC1 and also CA RACB recruited YFP-RIPa to the cell periphery, whereas DN RAC1 and DN RACB did not (Additional file 2).

**Fig. 3:**
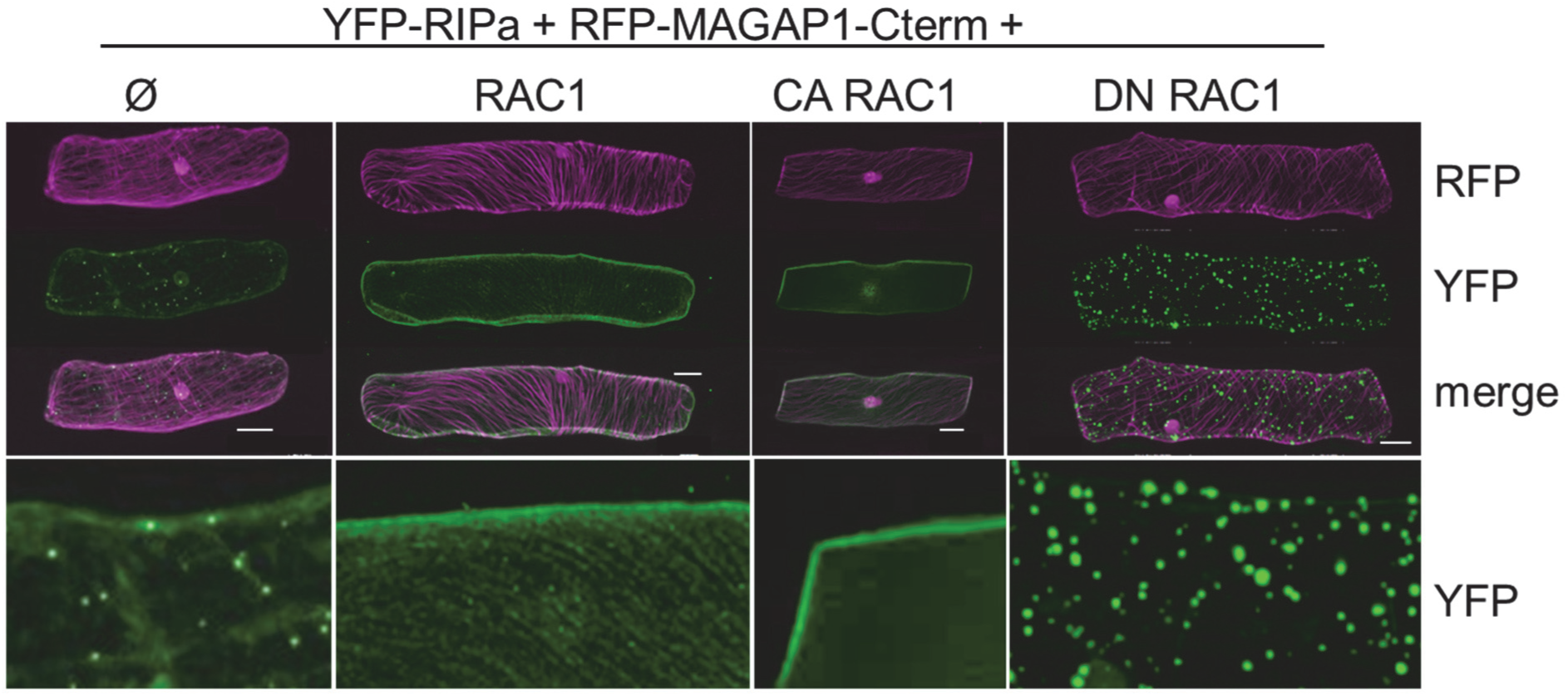
Single cell expressed barley YFP-RIPa changes subcellular localization upon co-expression of untagged RAC1. YFP-RIPa alone (Ø) localizes to immobile speckles and the cytoplasm. Co-expression of untagged RAC1 (WT RAC1) leads to plasma membrane and MT association of YFP-RIPs, co-expression of CA RAC1 leads to plasma membrane localization of YFP-RIPa and DN RAC1 leads to speckle-association of YFP-RIPa. The lower panels show digital magnifications of the YFP-RIPa signals with 40% enhanced brightness. Whole cell Z-stack images were taken 24 h after biolistic transformation of barley epidermal cells. The MT-marker RFP-MAGAP1-Cterm was co-expressed to visualize MTs. Bars represent 20 μm.

### A ROP – ROP-GAP module positions RIPa in MT-restricted domains at the cell periphery

Arabidopsis RIP3/MIDD1 localizes into MT-restricted membrane domains when co-expressed with the type II ROP ROP11, the catalytically active domain of ROP-GEF4 and ROP-GAP3 [16]. We hence speculated that co-expression of the barley ROP-GAP MAGAP1 and the barley type II ROP RAC1 could modulate subcellular localization of YFP-RIPa. Therefore, we first confirmed that MAGAP1 can interact with RAC1 in yeast and can recruit GFP-tagged MAGAP1 from MTs to the cell periphery/plasma memebrane (Additional file 3). We also found that MAGAP1 does not interact with RIPa in yeast (Additional file 1). We then used the MT marker DsRED-MAP4 and coexpressed it with YFP-RIPa, with untagged MAGAP1 and untagged RAC1. This led to accumulation of YFP-RIPa in MT-restricted domains at the cell periphery/plasma membrane. In this situation, MT-rich and YFP-RIPa-rich domains of the cell periphery mutually excluded or depleted each other (Figure 4). Similar images were recorded when we used RFP-MAGAP1-Cterm as an alternative MT marker. Additionally, MTs appeared to function in formation or restriction of the YFP-RIPa-enriched domains because treatment with 30 μM of the MT-depolymerizing drug oryzalin led to both disaapearance of detectable MTs and the destruction of these domains and to more evenly peripheral localization of YFP-RIPa (Figure 5). We also wanted to get more evidence for importance of MAGAP1 in heterogeneity of the YFP-RIPa distribution. Therefore, we co-expressed RAC1 and YFP-RIPa with different versions of labelled RFP-MAGAP1 to see whether a functional ROP-GAP is required to form the observed YFP-RIPa membrane domains. We used either full length RFP-MAGAP1 or a version, which lacked the carboxyterminal MT-assocciating domain (MAGAP1-ΔCterm), or the MT marker RFP-MAGAP1-Cterm, which lacks the ROP-binding CRIB and GAP domains (see figure Fig. 6A for domain composition of MAGAP1 versions). In these experiments we did not co-express untagged MAGAP1. Again, co-expression of full length RFP-MAGAP1 resulted in patchy domains of YFP-RIPa at the cell periphery/plasma membrane, which were restricted by RFP-MAGAP1 labelled MTs. Interstingly, using RFP-MAGAP1-Cterm instead of full length RFP-MAGAP1, completely dissolved the accumulation of YFP-RIPa in specific membrane domains but showed YFP-RIPa distribution at the entire cell periphery/plasma membrane. Hence, the ROP-interacting domains of MAGAP1 appeared to be necessary for the formation of distinct YFP-RIPa-labelled membrane domains. Strikingly, when we used RFP-MAGAP1-ΔCterm, this protein seemed to be recruited by RAC1 to the cell periphery/plasma membrane and YFP-RIPa appeared again in speckles of unknown nature. This suggests that RFP-MAGAP1-ΔCterm outcompeted YFP-RIPa from the interaction with RAC1 and hence a pattern occurred that is similar to that observed under co-expression of DN RAC1, which does not bind RIPa (compare Figs. 1 and 3).

**Fig. 4.**
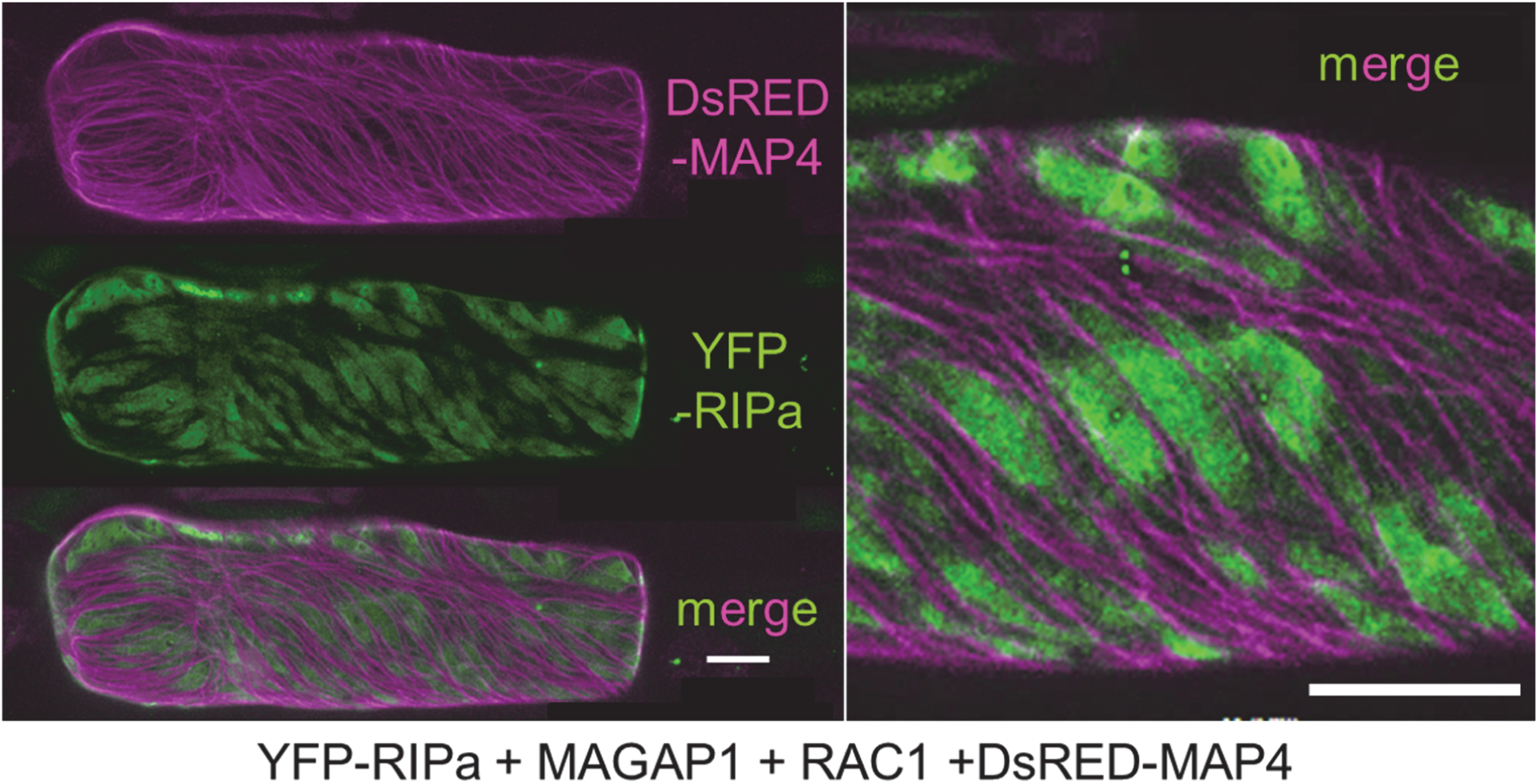
Barley YFP-RIPa localizes to MT-restricted domains of the cell periphery/plasma membrane when co-expressed with wild type RAC1 and MAGAP1. Whole cell Z-stack images were taken 24 h after biolistic transformation of barley epidermal cells. The MT-marker DsRED-MAP4 was co-expressed to visualize MTs. The right panel shows a digital magnification with 40% enhanced brightness. Bars represent 20 μm.

**Fig. 5.**
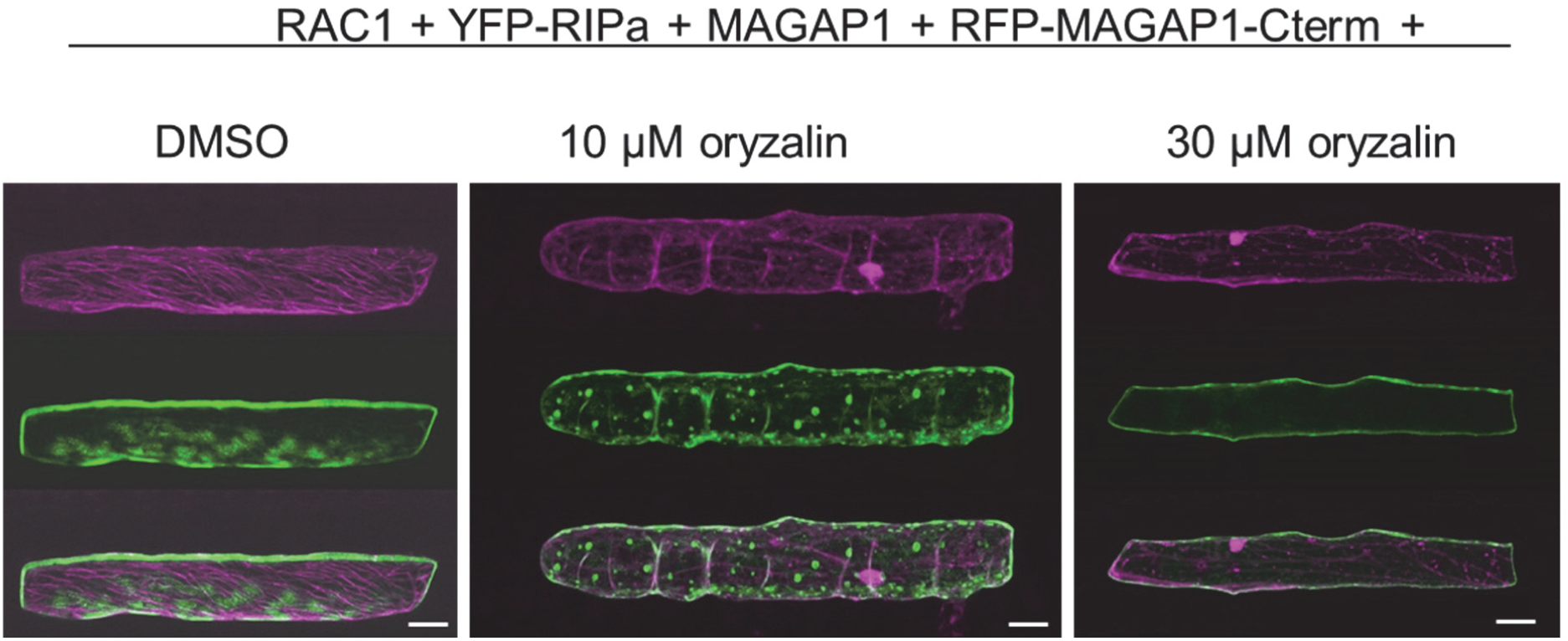
Disturbance of YFP-RIPa localization to MT-restricted domains of the cell periphery/plasma membrane. YFP-RIPa (shown in green) when co-expressed with wild type RAC1 and MAGAP1 can be found in MT-restricted domains of the cell periphery/plasma membrane (see left panel for DMSO solvent control). This localization is dissolved when MTs are destroyed by either 10 or 30 μM oryzalin treatment (solved in 0,25% [v/v] DMSO, treated for 3,2h before imaging). Whole cell Z-stack images were taken 24 h after biolistic transformation of barley epidermal cells. The MT-marker RFP-MAGAP1-Cterm (shown in magenta) was co-expressed to visualize MTs. Bars represent 30 μm.

**Fig. 6.**
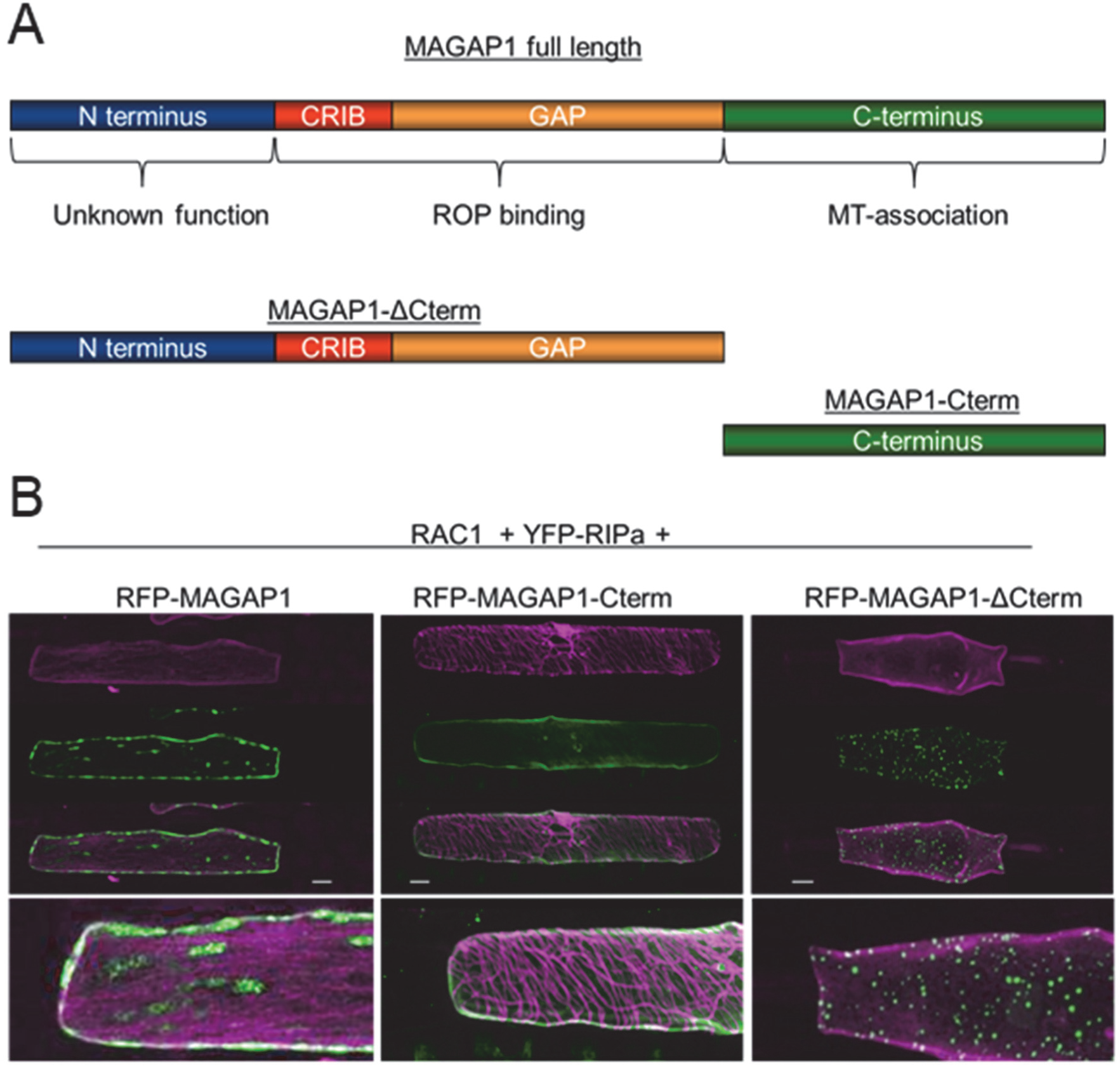
Functional domains of MAGAP1 determine the formation of YFP-RIPa membrane domains. A. Domain architecture of MAGAP1 and of truncated versions of MAGAP1, which were expressed as RFP fusion proteins. B. Co-expression of fluorescent YFP-RIPa (shown in green) and untagged RAC1 with three versions of RFP-MAGAP1 (shown in magenta) with or without ROP-binding and MT-association domains as depicted in A. Whole cell Z-stack images were taken 24 h after biolistic transformation of barley epidermal cells. Bars represent 20 μm.

### RIPa accumulates at sites of fungal attack

When transiently over-expressed in barley epidermal cells, CA RAC1 does not significantly support or inhibit penetration by *Bgh*. We also did not measure a significant influence of transient RIPa over-expression on *Bgh* penetration success, when we applied the exact experimental prtotocol, in which RIPb over expression supports fungal penetration [18]. Yeast-two-hybrid assays did not suggest a direct interaction between RIPa and the *Bgh* virulence effector ROPIP1, which may target barley RACB but can also bind RAC1 in yeast [11] (Additional file 1). We hence wondered how YFP-RIPa would localize in interaction with *Bgh*. When we inoculated leaves, in which we co-expressed YFP-RIPa, RAC1, MAGAP1 and the MT marker RFP-MAGAP1-Cterm, we detected, albeit somewhat less clear than in non-inoculated leaves, patterns of mutually exclusive MTs and YFP-RIPa-labelled membrane domains. Additionally, YFP-RIPa clearly labelled a zone around the site of fungal attack likely representing plasma membrane that directly attached to the defensive cell wall apposition that barley forms in response to the penetration attempt from the fungal appressorium (Fig. 7) [20]. Since we expressed RAC1 in its wild type form in these experiments, we also inoculated cells expressing YFP-RIPa under co-expression of CA RAC1 or DN RAC1. This revealed that YFP-RIPa localized to sites of fungal attack in cells with CA RAC1, too, but remained in unknown speckles, when co-expressed with DN RAC1 (Additional file 4).

**Fig. 7.**
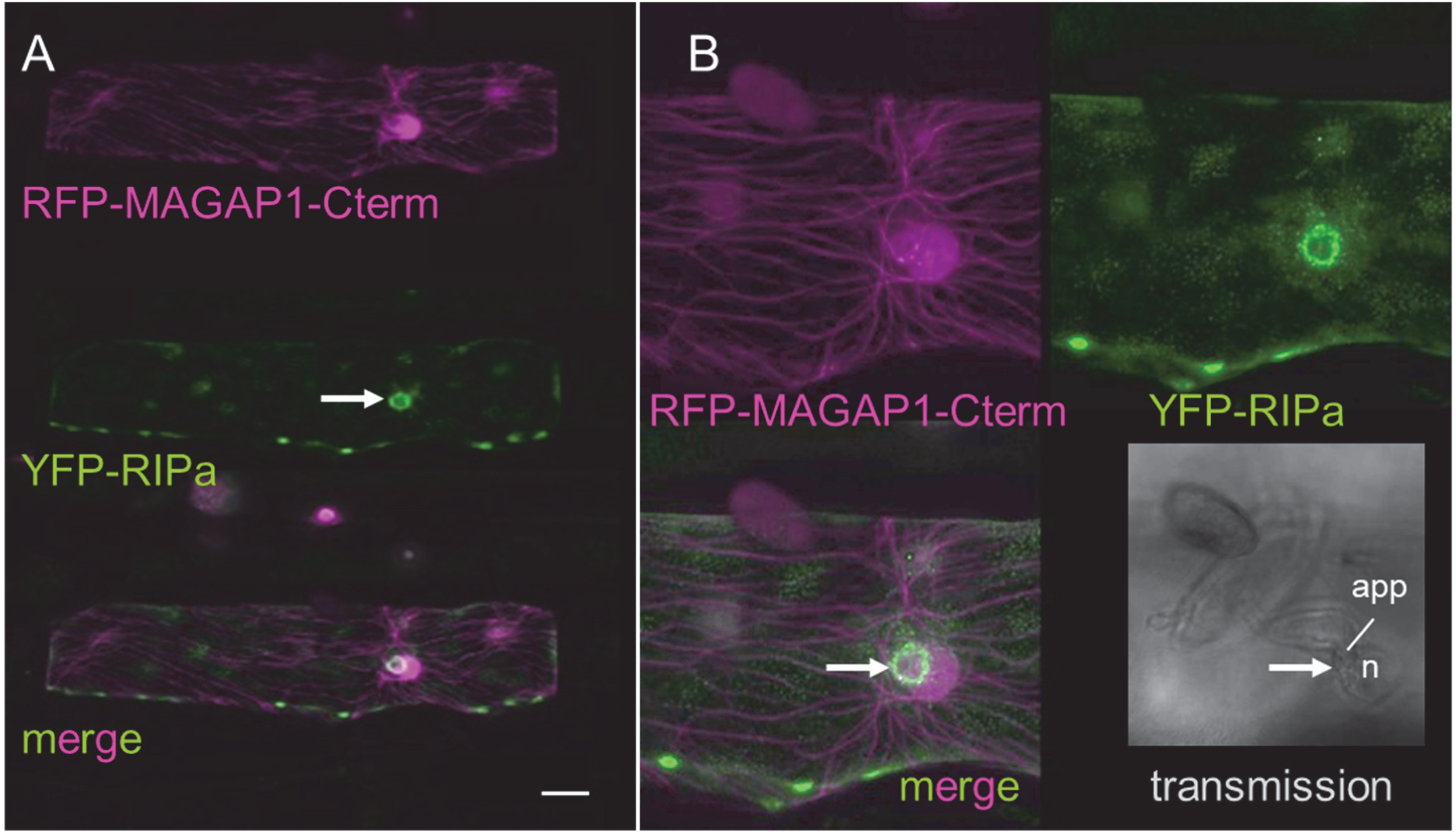
YFP-RIPa localization at sites of fungal attack by *Bgh*. A. Whole cell Z-stack images were taken 28 h after biolistic transformation of barley epidermal cells and 23 h after inoculation. The MT-marker RFP-MAGAP1-Cterm was co-expressed to visualize MTs. Bars represent 20 μm. Additionally, untagged RAC1 and untagged MAGAP1 are co-expressed. B. Same cell as in A imaged at a higher zoom factor. Brightness was enhanced by 40 % after imaging. Please note the fungal attack from an appressorium (app). YFP-RIPa is visible in membrane patches and around the site of attack (arrow). n, plant nucleus.

## Discussion

### RIPa is a ROP-binding protein

Signalling RHO GTPases are crucial for cell polarity and cell development across the border of kingdoms. In plants, ROPs are increasingly well understood as molecular hubs that integrate signals from the cell periphery or apoplast and hormone responses to translate this into cellular organization of the cytoskeleton or membrane trafficking machinery. This serves among others polar cell development or response to pathogens and cell wall sensing [4, 21, 22]. To translate signalling cues into downstream-signalling ROP-GTP interacts with so-called ROP-effectors that either perform a direct function or serve as scaffolds for recuitment of other downstream factors in higher order complexes. The knowledge on plant ROP-effectors is constantly increasing but still very incomplete and for many ROP-effectors, we lack knowledge about the molecular mechanism, by which they control cellular organization [3]. Therfore and because ROP signaling is involved in plant resistance and susceptibility to diseases, we are interested in finding further ROP-effectors. We search for them in barley, because i. in monocot crops the knowledge on ROP signalling is even less complete than in Arabidopsis, ii. barley ROPs are involved in pathogensis of powdery mildew, and iii. the interaction of plants with powdery mildew fungi is a model system for studying the cell biology of plant-microbe interactions [23]. Based on what we and others found for RIP/ICR proteins in Arabidopsis, we identified barley RIPa as a candidate ROP-effector. We found that it preferentially interacts with the activated form of both type I and type II ROPs. This is similar to RIPs of Arabidopsis, which interact with diverse ROPs in yeast. Additionally, there is also genetic interaction of ROPs and RIPs *in planta* [14, 24, 25] [17]. In addition to our yeast-based interaction assays, the dynamics of subcellular RIPa localization upon co-expression of different versions of ROPs suggest that ROPs can interact with RIPa *in planta*. The fact that constitutively GTP-loaded CA RAC1 and wild type RAC1, which can be naturally loaded with GTP, recruited RIPa to the cell periphery strongly supports that RIPa interacts with signalling forms of ROPs such as RAC1-GTP at the plasma membrane. The partial accumulation of RIPa in unknown speckles, when overexpressed alone or with DN RAC1 or DN RACB further suggests that RIPa without a matching amount of binding partner forms aggregates or accumulates in unidentified cellular compartments. This is different to barley RIPb, which we recently found in the cytosol, at MTs and the cell periphery, when expressed alone. However, RIPb is naturally expressed on a higher level in the barley epidermis, when compared to RIPa, and hence might be also co-expressed with higher amounts of natural binding partners in the barley epidermis [18].

### ROP activity and MTs control symmetry breaking of plasma membrane domains labelled by RIPa

The recuitment of RIPa by CA RAC1 or CA RACB suggested that the membrane association of RIPa depends on ROP signalling activity. We hence tested whether we can reconstitute a ROP-activation-deactivation module similar to what was reported for Arabidopsis xylem, in which RIP3/MIDD1 coordinates locally restricted cell wall apposition, and *Nicotiana benthamiana* epidermal cells. In these models, expression of ROP11, the catalytic domain of ROP-GEF4, ROP-GAP3 and RIP3/MIDD1 provokes symmetry breaking of the plasma membrane into zones with high and low ROP activity. This becomes visible by the presense of RIP3/MIDD1 in membrane domains of high ROP activity [16, 22]. RIP3/MIDD1 can further interact with kinesin13A *in planta* [14] and recruits this protein into areas of high ROP activity, where it supports the depolymeritzation of MTs from the plus end. *Vice versa*, MTs laterally restrict RIP3/MIDD1-labelled ROP activity domains leading to lateral mutual inhibition of MTs and ROP activity and depletion of MTs from zones of high ROP11 activity [15]. Interestingly, the expression of RAC1 and MAGAP1 together with RIPa appeared to be suffient to reconstitute a MT-controlled ROP-activation-deactivation module in barley. Asymmetric appearance of RIPa at the plasmamembrane in zones with very few or mostly lacking cortical MTs was reminiscent of the RIP3/MIDD1-labelled domains to ROP activity in Arabidopsis. We did not co-express any ROP-GEF in these cells and hence it seems that the barley epidermis possesses sufficient endogenous GEF activity to activate RAC1. This is further supported because expression of wild type RAC1 similar to expression of CA RAC1 recruited RIPa to the plasmamembrane in cells without co-expression of MAGAP1. We assume that RAC1 was activated by barley endogenous ROP-GEFs in these situations but hardly deactivated becaue no correspondingly high amount of ROP-GAP was present in those cells, and ROPs have only a weak intrinsic GTP-hydrolyzing activity [26]. However, additional co-expression of either untagged MAGAP1 or RFP-tagged MAGAP1 led to symmetry breaking of the plasma membrane. MAGAP1 may not directly interact with RIPa but with activated RAC1 in theses situations as our yeast-two-hybrid assays support. Hence, MAGAP1 might fulfil a complex function in these situations. One the one hand, MAGAP1 is a classical ROP-GAP with a CRIB domain that supports binding to ROP-GTP and possesses a conserved catalytical arginine, which is predicted to hydrolyze ROP-bound GTP and appears to be required for the control of ROP effects [9]. On the other hand, MAGAP1 is directly associated to MTs by its carboxyterminal domain and hence ideally positioned to perform a function in spatial feedback from MTs. This is different from Arabidopsis ROP-GAP3 for which no MT-association is reported. The idea, that MAGAP1 indeed function in lateral restriction of ROP activity domains in barley is strongly supported by the expression of truncated versions of MAGAP1, which interfered with membrane symmetry breaking. RIPa speckles were observed, when we co-expressed RAC1 with MAGAP1-ΔCterm, which is detached from MTs by truncation of its C-terminus but possesses intact domains for ROP-GTP interaction and GTP hydrolysis [9]. Catalytic activity of MAGAP1-ΔCterm is supported because it is fully functional in limiting susceptibility to *Bgh* [9]. In this situation, MAGAP1-ΔCterm occurred at the plasma membrane, to which it was most likely recruited by the co-expressed RAC1. We speculate that MAGAP1-ΔCterm outcompetes RIPa from binding to RAC1 in this situation and additionally functions as a ROP-GAP such that most of the expressed RAC1 is deactivated immidiately after loading GTP. Together, this could explain occurrence of RIPa in speckels, in which it otherwise was observed without co-expression of RAC1 or upon co-expression of DN RAC1. By contrast, RIPa more symmetrically labelled the cell periphery when MAGAP1-Cterm was expressed, which does not possess any ROP binding or GAP activity domain but still localizes to MTs. This also shows that MTs did not serve as a pure physical barrier to the diffusion of RIPa or RAC1 activity but as a physiological barrier dependent on a the presense of full length MAGAP1. Together, both GAP activiy and the spatial control of this activity near MTs appear nesesarry for symmetry breaking of ROP activity at the plasma membrane (see also Additional file 5 for a model). MAGAP1 has been suggested to function in MT-associated feedback on ROP activity in barley [9].

### RIPa might label a membrane domain of high ROP activity in interaction with Bgh

In *Bgh*-attacked cells, RIPa was also observed in membrane domains, when co-expressed with RAC1 and MAGAP1. However, the lateral restriction of RIPa-domains by MTs was less distinct. The overall intensity of RIPa labelling of the membrane was not very high when contrasted by local accumulation at the site of fungal infection. Because RIPa seems to prefentially accumulate at sites of high ROP or more specifically RAC1 activity, this might indicate that RAC1 can be activated at sites of fungal attack. This is reminiscent of the accumulation of further ROP activity sensors such as RIC171 or RIPb at sites of fungal attack [27][18]. Together, these observations support earlier hypotheses of locally enhanced ROP activity at sites where *Bgh* attempts to penetrate [27, 28].

The physiological effect of this local ROP activity is not well understood and RIPa has no significant effect on the fungal penetration success when over-[18]. RAC1 seems to be involved in modulation of fungal penetration success in barley but this depends on whether CA RAC1 was expressed transiently or stably and on whether *Bgh* or *M. orzae* was attacking [7, 12]. The putative rice ortholog of barley RAC1 is also called RAC1. Rice RAC1 functions in chitin-triggered immunity and is activated via the chitin-signalling receptor kinase CERK1 and RAC-GEF1, a member of a plant-specific RHO-GEF family [29]. Chitin is a potent elicitor of early defense reactions in barley and can induce systemic resistance to *Bgh* infection [10, 30]. However, it is unclear to what extent chitin elicitation contributes to basal resistance of barley in the authentic interaction with *Bgh*. We can only speculate that chitin elicitation is also involved in local activation of RAC1 in barley but this would explain why we observe local enrichment of the RAC1 activity sensor RIPa at sites where we can assume chitin elicitors from the fungal cell wall to be present.

## Conclusions

Data suggest that barley RIPa interacts with barley ROPs and specifies RAC1-activity associated membrane domains with potential signaling capacity. Lateral diffusion of this RAC1 signaling capacity is restricted by microtubules and MICROTUBULE-ASSOCIATED ROP-GTPASE ACTIVATING PROTEIN 1. Hence, an interplay of ROP activity and spatially confined MT-associated enzymatic restriction of ROP activity by MAGAP1 can provoke symmetry breaking at the plasma membrane of barley epidermal cells. Resulting membrane heterogeneity potentially reflects a mechanism by which monocot cells focus ROP activity comparable to what was reported before for dicots. Focal accumulation of RIPa at sites of fungal attack may further indicate locally restricted ROP activity at sites of fungal invasion.

## Methods

### Plant and fungal material

We used the barley (*Hordeum vulgare*) cultivar Golden Promise for transformation and inoculation experiments. We gew plants with a light dark cycle of 16h/8h at light intensity of 150 μM s^−1^ m^−2^ and 65% relative humidity and at 18°C. *Blumeria graminis* f.sp. *hordei* race A6 was maintained on Golden Promise plants under the same conditions inoculated on plants by skaking plants with sporulating powdery mildew and blowing spores into a plastic tewer (200×50×50cm), which we had postioned over the naïve plants or transformed leaf segements on agar plates.

### Construction of expression constructs

Barley *RIPa* (HORVU3Hr1G087430) was amplified from cDNA using gene-specific start to stop primers equipped with Xba1_fwd and Xba1_rev restrction sites for subcloning (RIPaXbaI_fw 5’-TCTAGATATGCAGACAGCCAAGACAAG-3’; RIPaXbaI_rv 5’-TCTAGATCATTTCTTCCACATTCCACTG-3’). We ligated the amplicons into the pGEM-T easy vector (Promega, Madison, WI, USA) by blunt end cloning according to the manufacturer’s instructions and sequenced the inserts. For Yeast Two-Hybrid assays *RIPa* was sucloned from the pGEM-T easy vector into pGADT7 plasmid (Clontech Laboratories) using the mentioned restriction sites. For over-expression and protein localization we used the high copy pGY1 plasmid, containing the CaMV35S promotor. We cut the RIPa insert by Xba1 from the pGEM-T easy vector and ligated *HvRIPa* into the pGY1 plasmid or pGY1-YFP (without YFP STOP codon) plasmid to gain a N-terminal YFP fusion construct pGY1-YFP-RIPa. Orientation was confirmed by sequencing. For cloning into the Y2H pGADT7 vector, RIPa was emplified with RIPa_Nde 5’-TGGATCCTCATTTCTTCCACATTCCACTG-3’ and RIPa_BamH1 5’-ACATATGCAGACAGCCAAGACAAGG-3’. Construction of plant expression and Y2H vectors for barley MAGAP1, RAC1 and RACB variants was described previously [7, 9, 27]. Also, the construction of MAGAP1, RFP-MAGAP1 and truncated versions of this was described previously [9].

### Biolistic transformation of barley leaf segments

We transformed barley epidermal cells by biolistic particle bombardment with PDS-1000/HE (Biorad, Hercules, CA; USA) as described earlier [31]. Therefore, we placed segments of 7d old primary leaves of barley on 0.8-1% (w/v) water-agar. For each shot, we precipitated 1μg plasmid DNA on 302.5 μg of 1μm gold particles (Biorad, Hercules, CA, USA)by adding the same volume of 1M CaCl_2_. Half the DNA amount was used for pGY1-mCherry transformation markers. Finally, we added 3μl per shot of 2mg/ml protamine (Sigma) were. We subsequently (30 min later at RT) washed twice the plasmid-coated gold with 500μl of first 70% (v/v) and second 100% ethanol. The resuspendend gold particle were then pipetted (6 μl) on the macro carrier for bombardment.

### Subcellular localization and protein recruitment in planta

Localization of YFP-HvRIPa either expressed alone or simultaneously with different vcerions of RAC1, RACB and MAGAP1 was performed at the indicated time points after transient transformation of barley leaves. We imaged single transformed cells with a Leica TCS SP5 confocal laser scanning microscope and the use of hybrid HyD detectors. Excitation and emission wavelength were individuall adapted to the respective fluorophores as described before and imaged were recorded by sequentially scanning line-by-line with a 3-times averaging [9, 27].

### Yeast two-hybrid assays

Constructs were transformed into yeast strain AH109 following the small-scale LiAc yeast transformation procedure from the Yeast Protocol Handbook (Clontech, Mountain View, CA, USA). Bait- and prey-construct transformed yeast cells were dropped on either transformation-selected (SD -L-W) or interaction-selective (SD –L,-W,-A-H) medium. pGADT7 and pGBKT7 were inclued as empty vector controls to exclude auto-activity of respective constructs.

## Abbreviations

*Bgh*: *Blumeria graminis* f.sp. *hordei*;
CA: constitutively activated;
CRIB: CDC42/RAC-Interactive Binding;
DN: dominant negative;
GAP: GTPase-activating protein;
GEF: guanine nucleotide exchange factors;
ICR: Interactor of Constitutive Active ROPs;
RAC: Ras (Rat *sarcoma*)-related C3 botulinum toxin substrate 1;
RIC: <u>R</u>OP-Interactive <u>C</u>RIB motif-containing proteins;
RIP: <u>R</u>OP-Interactive <u>P</u>artner;
ROP: RHO of plants
MIDD1: microtubule depletion domain 1;
MAGAP1: MICROTUBULE-ASSOCIATED ROP-GTPASE ACTIVATING PROTEIN 1;
MT: micotubule

## Ethics approval and consent to participate

Not applicable

## Consent for publication

Not applicable

## Availability of data and material

All the data supporting our findings is contained within the manuscript. Constructs and seeds are available upon request from TUM.

## Authors’ contributions

RH developed the research questions, designed the study, prepared figures and wrote the manuscript. CH and CM designed and performed the experiments and prepared figures.

## Acknowledgements

We are grateful to Stefan Engelhardt (TU Munich, Chair of Phytopathology) for technical advice and fruitful discussions and to Vera Schnepf (TU Munich Phytopathology) who performed pilot experiments on barley RIPa.

## ADDITIONAL FILES

**Additional file 1: Figure S1.**
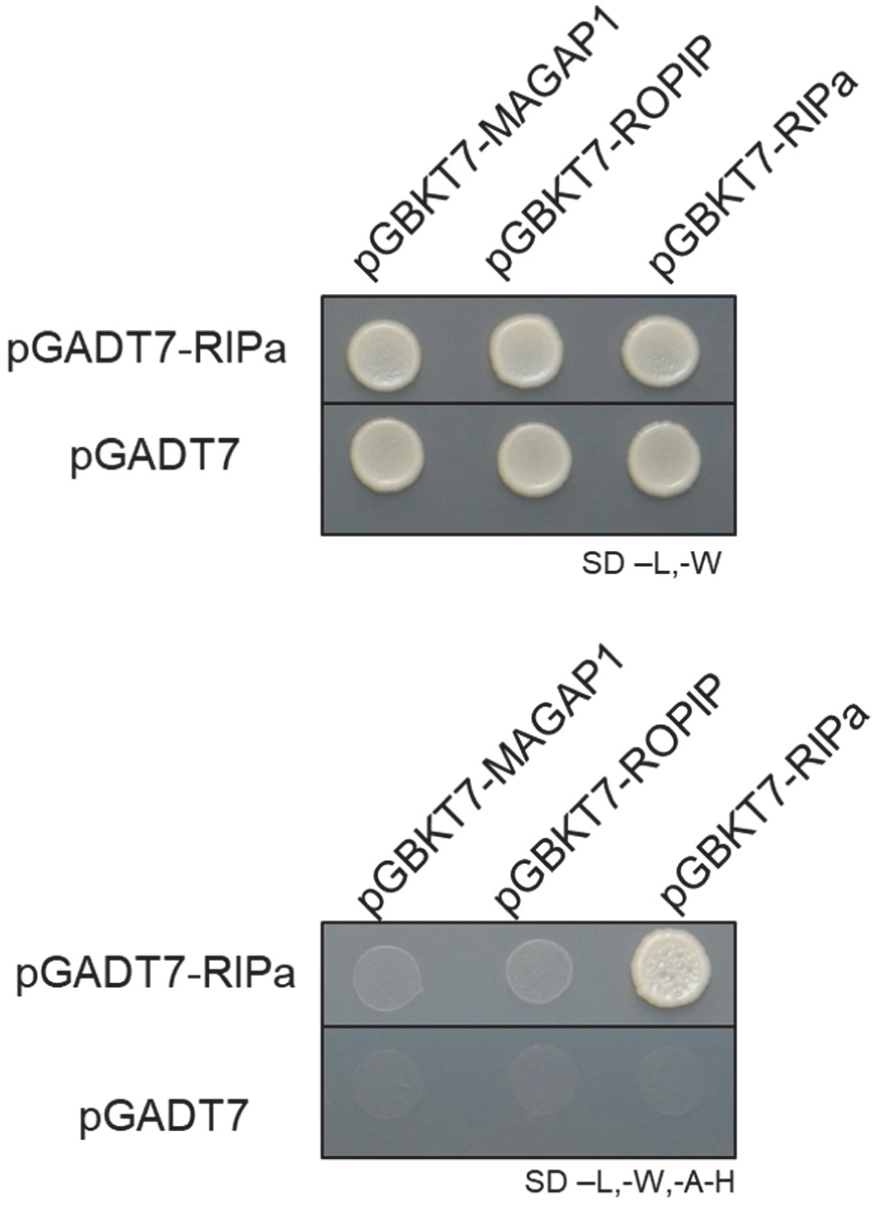
Barley RIPa interacts with itself in yeast. Bait- and prey-construct transformed yeast cells were dropped on either transformation-selected (SD -L-W) or interaction-selective (SD –L,-W,-A-H) medium. pGADT7 presents empty vector controls to exclude auto-activity of respective constructs.

**Additional file 2: Figure S2.**
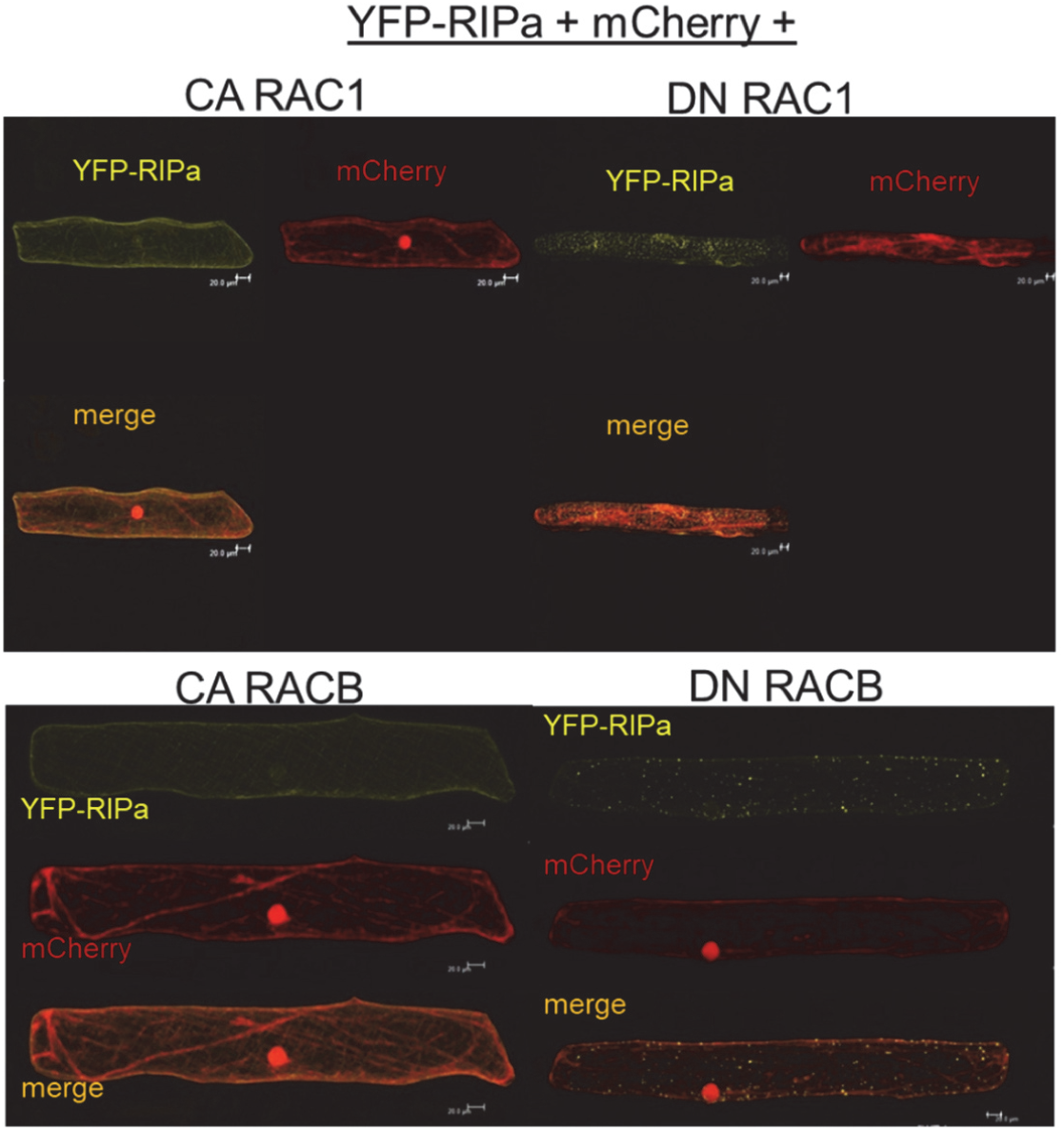
Barley YFP-RIPa localizes to the cell periphery when co-expressed with CA RAC1 or CA RACB (left panels) and to speckles of unknown nature when co-expressed with DN RAC1 or DN RACB (right panels. Whole cell Z-stack images were taken 24 h after biolistic transformation of barley epidermal cells. The cytosolic marker mCherry was co-expressed to contrast the cytoplasm.

**Additional file 3: Figure S3.**
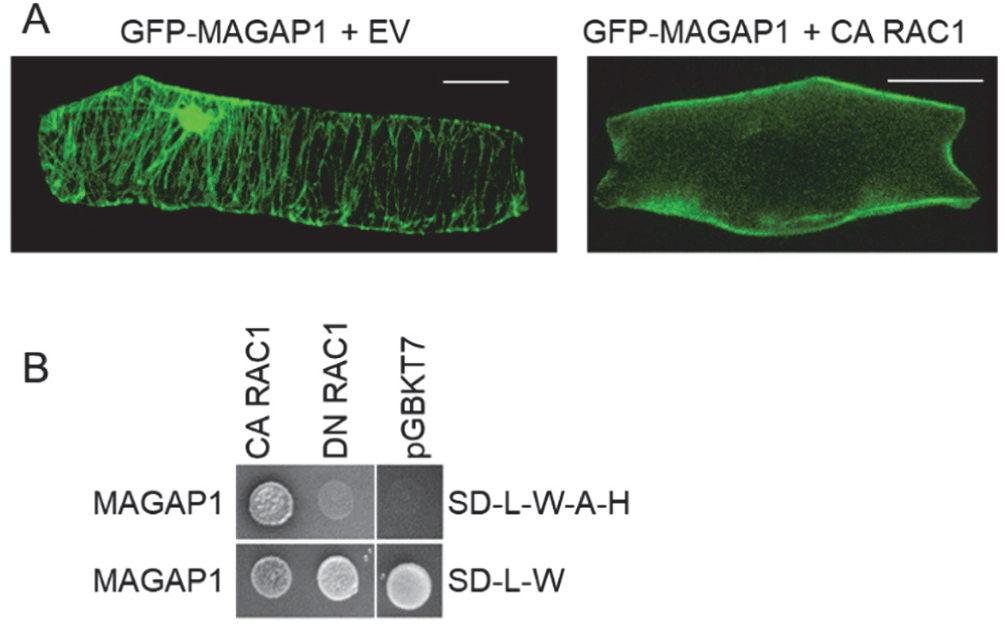
Interaction between MAGAP1 and RAC1. A. Change of GFP-MAGAP1 localization upon co-expression of CA RAC1 Whole cell Z-stack images were taken 24 h after biolistic transformation of barley epidermal cells. The MT-marker DsRED-MAP4 was co-expressed to visualize MTs. Bars represent 20 μm. B. C. Barley MAGAP1 interacts with the barley type II ROP RAC1 in yeast. Bait- and prey construct-transformed yeast cells were dropped on either transformation-selected (SD -L-W) or interaction-selective (SD –L,-W,-A-H) medium. pGADT7 and pGBKT7 represent an empty vector control to exclude auto-activity of the MAGAP1 construct.

**Additional file 4: Figure S4.**
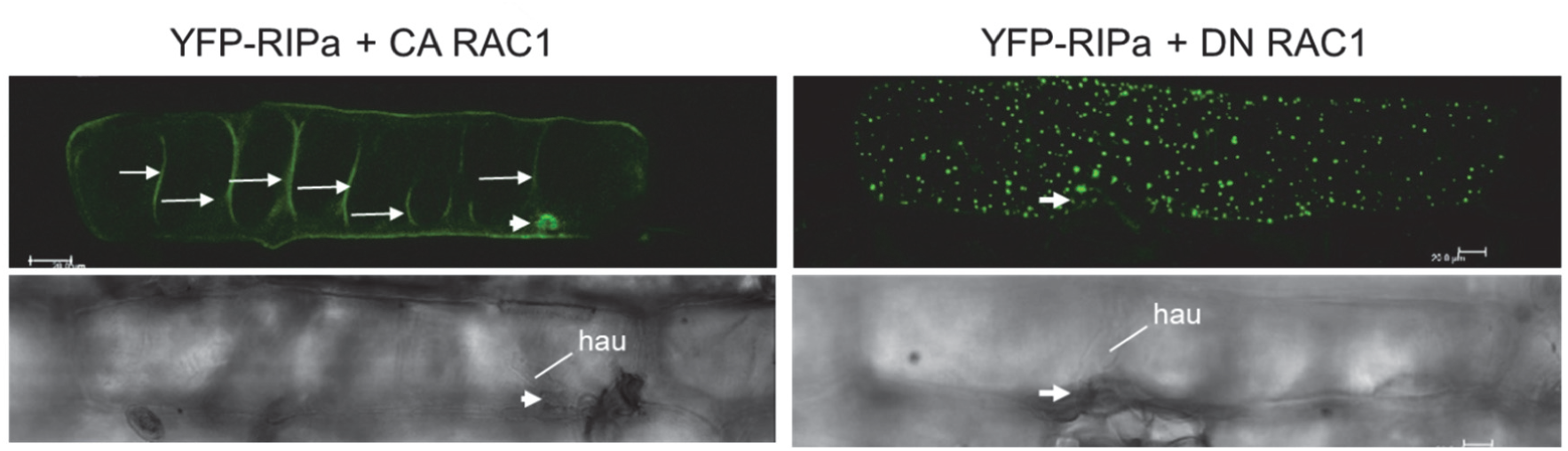
YFP-RIPa localization at sites of fungal attack by *Bgh* but not when DN RAC1 is co-expressed. Whole cell Z-stack images were taken 28 h after biolistic transformation of barley epidermal cells and 23 h after inoculation. Additionally, untagged CA RAC1 or DN RAC1 are co-expressed. Brightness was enhanced by 20 % after imaging. Please note the fungal attack from an appressorium (app). Site of attack, arrow; hau; fungal haustorium. Long arrows mark plasma membrane folds at cell wall protrusions at the cell bottom facing mesophyll cells. Bars represent 20 μm.

**Additional file 5: Figure S5.**
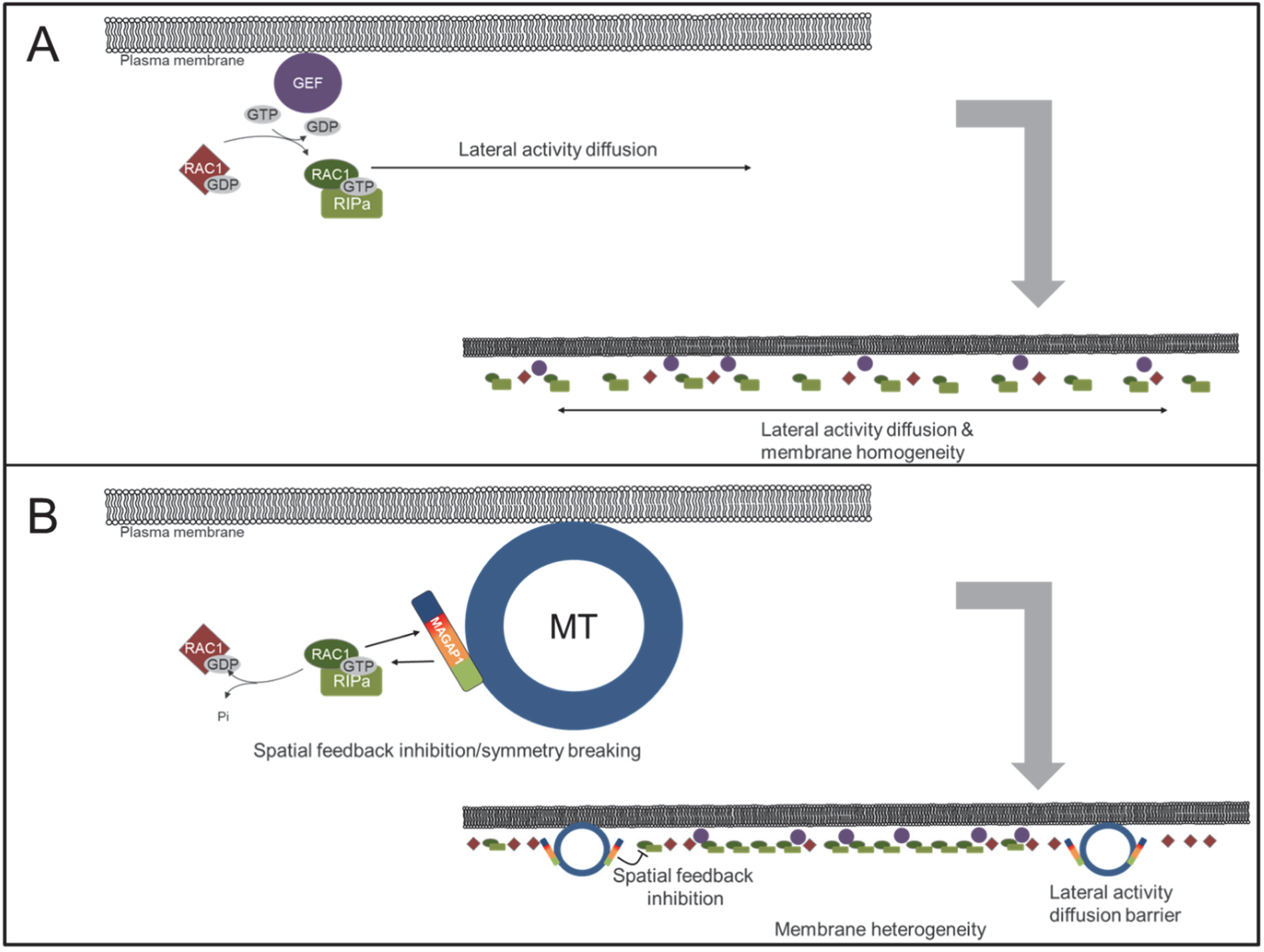
MT and MAGAP1-dependent symmetry breaking of plasma membrane-associated RAC1-RIPa signaling. A. In absence of MTs and MAGAP1, GEF-supported RAC1 activity can freely diffuse at the plasma membrane and RIPa is evenly distributed. B. In presence of intact MTs and functional MT-associated MAGAP1, MAGAP1 laterally inhibits RAC1 activity from MTs. This leads to spatially restricted negative feedback, and hence symmetry breaking and membrane heterogeneity.

